# Exploring Chromosomal Position Effects for Predictable Tuning of Metabolic Pathways in Yeast

**DOI:** 10.64898/2026.04.06.716637

**Authors:** Haosen Hong, Yuxin Cai, Jian Wang, Chang Dong, Qiang Zhang, Jiazhang Lian

**Affiliations:** Key Laboratory of Biomass Chemical Engineering of Ministry of Education & State Key Laboratory of Biobased Transportation Fuel Technology, College of Chemical and Biological Engineering, Zhejiang University, Hangzhou 310027, China; Zhejiang Key Laboratory of Intelligent Manufacturing for Functional Chemicals, ZJU-Hangzhou Global Scientific and Technological Innovation Center, Zhejiang University, Hangzhou 311200, China; ZJU-UIUC Institute, International Campus, Zhejiang University, Haining, Zhejiang, 314400, China; State Key Laboratory of Coordination Chemistry, School of Chemistry and Chemical Engineering, Nanjing University, Nanjing 210093, China

## Abstract

Precise control of gene expression is essential for optimizing metabolic pathways, yet current tuning strategies based on promoter strength or gene copy number remain largely empirical. Chromosomal position represents an additional regulatory axis, as identical gene expression cassettes can exhibit markedly different expression levels depending on their integration sites. Here, we systematically quantified the expression output of 98 intergenic regions (IGRs) in *Saccharomyces cerevisiae* using a fluorescent reporter and developed a predictive framework, Yeast IGR Prophet (YeIP), that infers expression potential directly from genomic context. By integrating multi-scale genomic features including transcriptional neighborhood, chromatin state, and chromosome topology, YeIP accurately predicted expression ranks and enabled the construction of a genome-wide atlas of expression hotspot and coldspot. Using this atlas, we rationally optimized a three-gene lycopene pathway solely through genomic integration site selection, achieving optimal transcriptional stoichiometry without modifying promoters or gene copy numbers. These results transform chromosomal integration sites from static genomic coordinates into programmable regulatory elements, establishing a predictive, data-driven framework for rational and scalable design of metabolic pathways in yeast.

## Introduction

The optimization of metabolic pathways is a central goal of synthetic biology, aiming to maximize the production of valuable compounds (1–3). Lifting up the expression level of rate-limiting enzymes by utilizing strong promoters or increasing gene copies usually boosts the yield of pathway products. However, naively maximizing the expression of all enzymes simultaneously often leads to suboptimal performance due to metabolic burden and the wasteful allocation of cellular resources to unnecessary protein production (4–6). This is because high expression of one enzyme can lead to the accumulation of toxic intermediates or the rapid depletion of essential precursors, creating metabolic bottlenecks that reduce the final yield. Consequently, achieving a precisely balanced transcriptional stoichiometry, where each enzyme abundance is carefully titrated, is now recognized as a more effective strategy for maximizing productivity. Classical strategies rely primarily on promoter tuning, copy number adjusting, and untranslated region (UTR) engineering to modulate transcriptional/translational outputs (7–9).

Chromosomal integration into intergenic regions (IGRs) introduces an additional, often underappreciated, regulatory dimension. The expression of an identical gene expression cassette can vary dramatically depending solely on its genomic location, a phenomenon known as the position effect (10). This variability arises from differences in local chromatin environment and neighborhood gene activity, has traditionally been viewed as a source of noise, and has been consistently reported in genome-wide studies (11–14). However, these findings have limited applicability to metabolic engineering, as most previous analysis involved replacing endogenous genes in single-knockout strains rather than integrating heterologous constructs into IGRs in common chassis organisms. Consequently, metabolic engineers have largely prioritized to identify a few “safe harbors” that support efficient and stable expression (15–19). Although recent studies have expanded the catalog of such sites to over 70 loci (20), this stability-oriented approach neglects a broader opportunity: the intrinsic variability of genomic loci can be harnessed as a tunable resource for pathway optimization. Rather than treating position effects as noise to be eliminated, they can be reframed as a continuous regulatory spectrum to be exploited. If the influence of a given IGR on gene expression could be predicted accurately, chromosomal position would become a programmable design parameter. By understanding this relationship, IGRs function as a novel regulatory element, enabling precise, context-dependent modulation of expression simply by selecting the appropriate integration site.

Here, we systematically quantified the expression output of 98 distinct IGRs in *Saccharomyces cerevisiae* using a constitutive fluorescent reporter to build a map of chromosomal positional effects directly relevant to pathway engineering. We then performed multi-scale feature engineering to encode the genomic context of each site, incorporating transcriptional neighborhood, epigenetic state, and higher-order chromosomal structure. These features enabled training of a machine learning model, Yeast IGR Prophet (YeIP), that predicts expression potential from genomic context alone and generalizes to unseen loci. This predictive capability transforms IGR selection from ad-hoc empirical screening into a rational pre-selectable design variable. Finally, we applied this framework to a three-step lycopene pathway to test whether genomic location selection alone can drive lycopene production tuning. Instead of maximizing the expression of all enzymes simultaneously, the most productive strain emerged from a balanced configuration resulted from strategic IGR allocation, without altering promoters, copy numbers, or coding sequences. We therefore establish IGR as a fine-tuning dimension that complements traditional promoter and UTR engineering. These results demonstrate that chromosomal integration sites can function as continuously tunable genomic rheostats, enabling predictable control over multi-gene expression ratios and establishing chromosomal position effects as a programmable engineering degree of freedom for synthetic biology.

## Materials and methods

### Strains and media

The *S. cerevisiae* strain BY4741 (*MATa; his3Δ1; leu2Δ0; met15Δ0; ura3Δ0*) was used as the host for all genetic modifications and phenotypic assays. The yeast strains harboring the reporter expression cassette were routinely cultivated in YPD medium or synthetic complete dextrose (SCD) medium, supplemented with the appropriate dropout nutrients as required. SCD-LEU-URA medium consists of 1.7 g/L yeast nitrogen base without amino acids and ammonium sulfate, 5 g/L ammonium sulfate, 20 g/L glucose, and a complete supplement mixture lacking leucine and uracil (CSM-LEU-URA), with 20 g/L agar added for solid medium. For testing in different carbon sources, SC medium was supplemented with 2% of either dextrose, galactose, raffinose, or glycerol. Cells were grown at 30°C with shaking at 800 rpm in 96-deep-well plates for 48 h to reach steady-state expression before flow cytometry analysis. *Escherichia coli* DH5α was employed for plasmid construction and propagation, and cultured in Lysogeny Broth (LB) medium at 37°C with shaking at 220 rpm. Ampicillin was added to a final concentration of 50 μg/mL for selection.

### Plasmid construction

The p415-iCas9 and p426*-SpSgH vectors were utilized for Cas9 expression and guide RNA (gRNA) cloning, respectively. Target-specific gRNA sequences were designed using Benchling (https://www.benchling.com), and the corresponding primers are provided in Supplementary Table S1 and S2. Following annealing and phosphorylation, the primer pairs were ligated into the p426*-SpSgH backbone pre-digested with BsaI-HFv2. The resulting constructs were transformed into *E. coli* DH5α, and correct insertion of the gRNA cassette was verified by DNA sequencing.

### Yeast transformation and genomic integration

Donor DNA fragments were generated via PCR amplification that using primers containing 40 bp homology arms flanking the specific integration site. For the fluorescent reporter, the *TDH3p-mCherry-ADH1t* expression cassette served as the template. Similarly, the carotenoid biosynthesis genes were cloned to construct donor fragments for lycopene pathway integration. Specifically, the *crtB* gene used in this study refers to a mutated variant of the *Xanthophyllomyces dendrorhous crtYB* gene (21), in which the lycopene cyclase activity was abolished to ensure exclusive lycopene accumulation. All the fluorescent reporter sequence used in this study are listed in Supplementary Table S3. Yeast transformations were performed using a modified lithium acetate/salmon sperm DNA/polyethylene glycol (LiAc/ssDNA/PEG) method. Mid-log phase BY4741 cells harboring the p415-iCas9 plasmid were rendered competent and co-transformed with 1 μg of the relevant gRNA plasmid and 1 μg of donor DNA. Heat shock was applied at 42°C for 1 h, after which transformants were selected on SCD-LEU-URA agar plates.

### Evaluation of integration efficiency and product yield

Individual transformants were inoculated into SCD-LEU-URA medium in 96-deep-well plates and cultured at 30°C with shaking at 800 rpm for 24 h. Cultures were then diluted for an additional 24 h. Fluorescence intensity was quantified using flow cytometry. Integration efficiency was calculated as the percentage of fluorescent-positive clones among eight parallel transformants screened per target site. For lycopene pathway integration, successful integrations were identified by Sanger sequencing and functionally verified by the concomitant development of a red colony phenotype, a visual trait conferred by lycopene accumulation. For phenotypic analysis of colony pigmentation, engineered strains were spotted onto SCD-LEU-URA agar plates and incubated at 30°C for 3 days before imaging. The strains carrying the *GAL1* promoter were grown to saturation in SCD medium, followed by subculturing at a 5% inoculum ratio into SC medium supplemented with 2% galactose. After induction for 24 h, the fluorescence intensity was determined.

Flow cytometry was performed on a BD LSRFortessa (BD Biosciences). mCherry fluorescence was excited with a 561 nm laser and analyzed using a 610/20 nm channel. For each sample, 30000 events were acquired. Cells were gated by FSC-A/SSC-A and FSC-A/FSC-H to exclude debris and aggregates, and fluorescence was quantified as mean intensity. Data were processed using FlowJo v10.8 (BD Biosciences).

A 200 µL aliquot of each culture was transferred to a 2 mL microcentrifuge tube, pelleted by centrifugation, and the supernatant removed. Cell pellets were resuspended in 1 mL DMSO and disrupted using a high-speed homogenizer to extract intracellular carotenoids. The clarified lysate was measured at OD_470_ to estimate lycopene content, while cell density was determined from OD_600_ measurements. Lycopene yield was expressed as OD_470_ normalized to OD_600_ to account for variations in biomass.

### RNA extraction and quantitative real-time PCR analysis

Total RNA was extracted from yeast cell pellets using the RNA Extraction Kit (Cat. HKR022, Hangzhou Haoke Biotechnology Co., Ltd., China) following the manufacturer’s protocol based on the Trizol-chloroform method. RNA concentration and purity were assessed using a CANHELP-100 ultramicro spectrophotometer (Canhelp Genomics). To eliminate genomic DNA contamination, RNA samples were treated with dsDNase. Subsequently, 1 μg of total RNA was reverse-transcribed into cDNA using the All-in-One First-Strand cDNA Synthesis SuperMix for qPCR (Cat. HKR041, Hangzhou Haoke Biotechnology).

Quantitative real-time PCR was performed on an ABI 7500 Real-Time PCR System (Applied Biosystems) using the 2X SYBR Green qPCR Master Mix (Cat. HKR040, Hangzhou Haoke Biotechnology). The 20 μL reaction mixture contained 10 μL of the master mix, 0.2 μM of both forward and reverse primers, and 1 μL of the cDNA template. The thermocycling program consisted of an initial pre-denaturation at 94°C for 2 min, followed by 50 cycles of 94°C for 5 s, 58°C for 10 s, and 72°C for 15 s, concluding with a standard melt curve analysis (60°C to 95°C). Relative gene expression levels were calculated using the 2^−ΔΔCt^ method. Target gene expression was normalized to the internal reference gene *ALG9* (ΔCt) and ΔΔCt values were computed relative to the designated calibrator for each experiment (specified in the corresponding figure legends; e.g., IntTrain92 for the promoter×locus panel and Strain D for the lycopene panel) and fold changes were reported as 2^−ΔΔCt^.

### Feature engineering

The workflow for establishing the training and prediction datasets is summarized in Supplementary Fig. S1. The 98 IGRs used for model training were selected based on high experimental integration efficiency (>75%) from our preliminary screens (22). IGRs were extracted from SGD R64-4-1 NotFeature FASTA (23). Integration sites were mapped onto IGR intervals, with interval length subsequently incorporated as a feature in YeIP. Autonomously replicating sequence (ARS) annotations from R64-4-1 were matched by chromosome. For each integration event, the distances to the nearest upstream ARS and downstream ARS elements were calculated, and the smaller of the two distances was recorded as the ARS distance feature. The transcriptional activity of the neighborhood was characterized using published cap analysis of gene expression (CAGE) sequencing data (24). Their annotated dominant transcription start sites (TSSs), which were defined based on the highest CAGE tag counts (TPM) within transcription clusters, were directly utilized. For each IGR, the dominant TSS located within a ±2 kb flanking window was mapped, and its reported TPM value was used as the “Neighbor Expression” feature. Essentiality was quantified by summing binary essential flags across a ±10 open reading frame (ORF) neighborhood flanking each integration site on the same chromosome (14). Nucleosome occupancy was assessed by summing measured nucleosomes signals within an inclusive ±0.5 kb window surrounding each integration site, representing local nucleosome density (25). Histone modification levels for H3K4me1 and H3K4me2 marks were computed by overlap-weighted averaging of bedGraph tiles strictly spanning each IGR interval (26). Chromatin boundaries and their associated strengths were obtained from a previous study (27). The boundary distance feature was calculated as the minimal distance between the integration site and the nearest chromatin block edges, while block strength corresponded to the measured strength value of the chromatin block containing the integration site. Finally, genome compact score at each integration site was quantified as the average compactness score of its flanking genes (27). To compare genomic features with different units on a common scale for visualization, feature-wise min–max normalization was applied to all features between the Top 20 and Tail 20 loci.

### Model training

A tabular regression model was trained to predict fluorescence intensity using AutoGluon-Tabular with the ’best’ quality preset (28). For model training and prediction, features were processed using AutoGluon’s default internal normalization pipeline. The input features were intergenic length, ARS distance, neighbor expression, neighbor essentiality, block strength, genome compact score, H3K4me2, H3K4me1, and nucleosome density. The dataset was split into 80% training and 20% test sets using a fixed seed. To curb overfitting, 5-fold bagging and one stacking level were used, which provides out-of-fold validation for model selection and blending. AutoGluon’s internal early stopping and missing-value handling were left at their default settings. The model was configured with problem_type = “regression”, num_bag_folds = 5, num_bag_sets = 1, and num_stack_levels = 1. Bayesian hyperparameter optimization was performed with a local scheduler for 50 trials. An ensemble model was then trained by AutoGluon, and a level-2 ensemble candidate was produced from the 5-fold cross-validation predictions; the best model or ensemble as determined by internal validation was used for inference on the held-out test set. Given the modest sample size, the Spearman rank correlation coefficient (SPCC) was prioritized to ensure the model has sufficient ranking ability for downstream IGR screening. SPCC was chosen as the primary metric as it assesses the ranking fidelity between predicted and observed values, which is critical for prioritizing engineering candidates, and is robust to non-normal distributions common in biological datasets. Mean squared error (MSE), coefficient of determination (R²), and pearson correlation (PCC) are additionally reported as reference metrics.

### Feature ablation analysis

Features were categorized into biologically distinct groups, where stand-alone models were trained exclusively. Models for ablation analysis were trained on this partial dataset, and its performance was evaluated on the test set.

### Statistical analysis and visualization

Data analysis was performed using Python 3.8. Statistical tests (Mann-Whitney U test, Spearman correlation) were calculated using SciPy package (v1.7.3). Figures were generated using Matplotlib package (v3.5.1) and Seaborn package (v0.11.2).

### Model interpretation using SHAP

SHapley Additive exPlanations (SHAP), a game-theoretic approach that assigns a contribution value to each feature, was employed to interpret the ensemble model. The model-agnostic KernelExplainer was utilized with a background dataset of 50 randomly sampled instances from the training set to estimate the expected model output. SHAP values were computed for the held-out test set to quantify the marginal contribution of each genomic feature to the predicted expression.

## Results

### IGRs feature engineering

Genomic integration of heterologous genes is one of the most widely used method to construct functional and genetical stable strains, which requires numerous neutral IGRs. In practice, the selection from IGRs usually follows 3 filters (18,29–35): (i) length should be exceeding 800 bp to minimize the potentially detrimental effects on the neighboring genes, (ii) IGRs should be far from the centromeres and telomeres, (iii) IGRs should be possessing high integration efficiency and minimal impact on cell fitness and genetic stability. To convert chromosomal context into a quantitative regulatory dimension, we first established a dataset linking genomic IGR context to heterologous expression output. We integrated a constitutive mCherry cassette into 98 distinct IGRs across the yeast genome, including several IGRs previously reported (20,22), measured steady-state fluorescence to quantify site-specific expression level (Fig. 1A), providing the training output for subsequent context feature engineering (Fig. 1B). We selected these IGRs within chromosomal arms, explicitly excluding regions within 0.5 kb of a centromere and 5 kb of a telomere to ensure genetic stability (Fig. 1C). We tested and compared all IGRs with IntTrain92 (hereafter referred to as “IntTrain loci”; Supplementary Table S1), a stable integration site showing the highest expression level in our previous study (22). Resulted fluorescent intensity exhibits a broad, unimodal range across loci (Fig. 1D; Supplementary Table S4), consistent with the previous reported strong position effects (20). The length distribution of IGRs is right-skewed, while centered around ∼1 kb, the length of several long IGRs reaches ∼4 kb, matching the engineering heuristic to prefer longer intergenic spaces (Fig. 1E).

**Figure 1.**
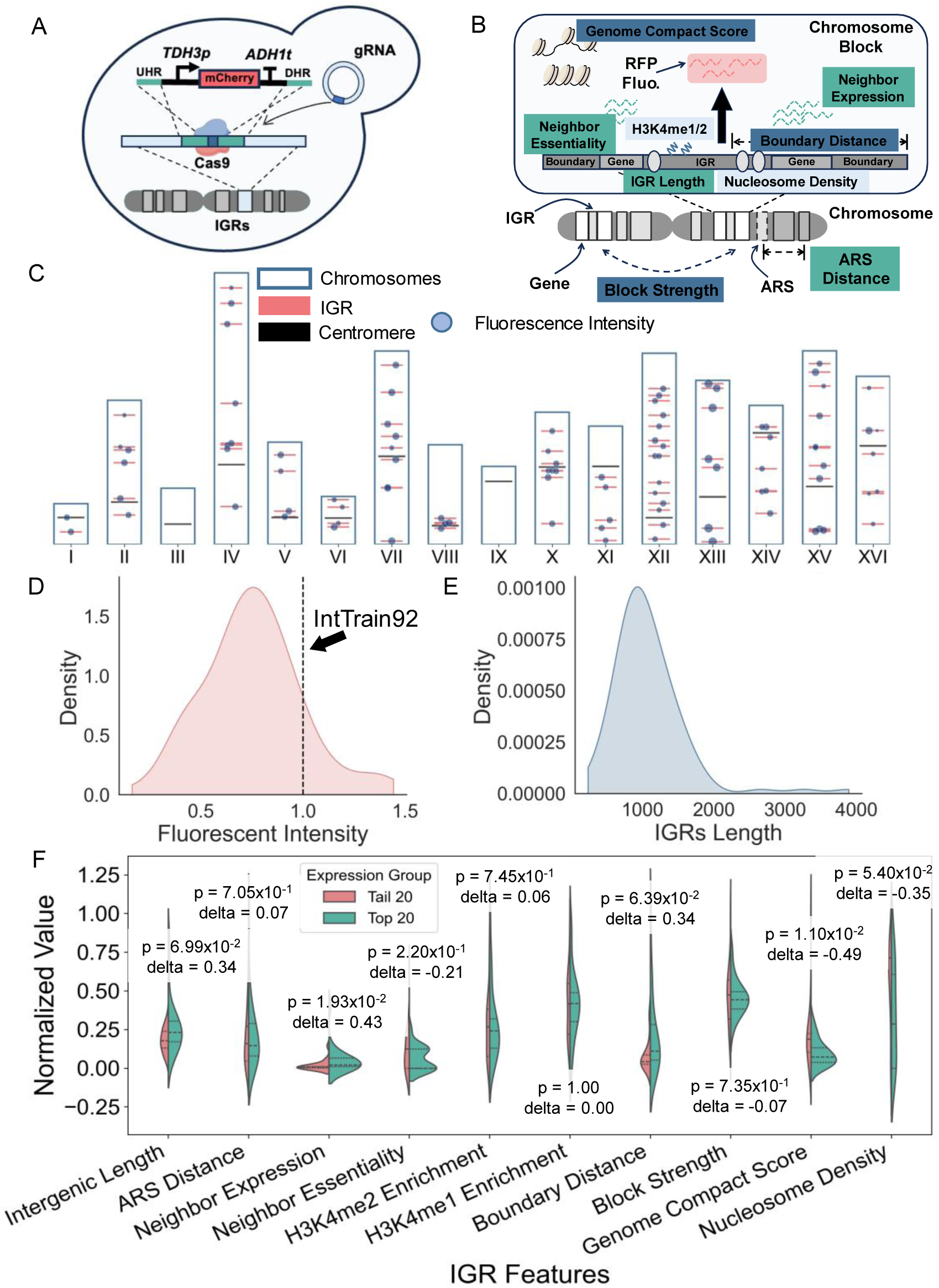
Experimental design, dataset characterization, and feature analysis for YeIP. (A) Schematic of the experimental strategy for characterizing intergenic regions (IGRs). A reporter cassette expressing mCherry from the constitutive *TDH3* promoter was integrated into 98 distinct IGRs in *S. cerevisiae* using a CRISPR-Cas9 system. (B) Schematic illustrating the features engineered to describe the genomic context of each IGR. Features are color-coded by category: Transcriptional Neighborhood (Green), Epigenetic Profile (Light Blue), and Chromatin Structure (Dark Blue). (C) Chromosomal distribution of the 98 characterized integration sites. Each blue dot represents an individual site, with the circle size proportional to the measured fluorescence intensity. Red lines indicate the mean fluorescence intensity for all sites on that chromosome, and gray bars denote the centromeres. (D) Density plot showing the distribution of measured mCherry fluorescence intensity. The values are normalized with IntTrain92 (*TDH3p*, SCD medium) indicated by the vertical dashed line (Supplementary Table S1). (E) Density plot showing the distribution of the physical length of the IGRs across the dataset. (F) Violin plots comparing the normalized feature distributions for the 20 IGRs with the highest expression (Top 20, teal) against the 20 with the lowest expression (Tail 20, red). P-values from a Mann-Whitney U two-sided test and the effect size (Δ) are shown.

To quantify and predict how genomic location impacts heterologous gene expression, we performed systematic feature engineering to describe the position effects of IGRs (Fig. 1B). Inspired by prior studies on chromosomal context (see methods), we organized these features across three dimensions. The first dimension captured transcriptional neighborhood composition and resource availability, represented by intergenic length, distance to ARS elements, local gene expression output, and essential gene density. The second dimension reflected the local epigenetic state, characterized by H3K4me1/H3K4me2 enrichment and nucleosome density. The third dimension represented higher-order chromosome structural organization, quantified using chromatin boundary proximity, chromatin block strength, and genome compactness. We designed such a multi-scale approach to provide a comprehensive description of the regulatory environment of each IGR.

To evaluate the effectiveness of the feature engineering, we compared feature distributions between the 20 highest-expressing and 20 lowest-expressing IGRs (Fig. 1F). Multiple features exhibited clear directional shifts, supporting their relevance. We observed the most pronounced separation for the Genome Compact Score, which was significantly lower in the high-expression group (p = 1.10×10⁻², Δ = -0.49), consistent with the principle that more accessible chromatin environments promote transcription. Neighbor Expression was also significantly higher in the high-expression IGRs (p = 1.93×10⁻², Δ = +0.43), indicating that transcriptionally active neighborhood facilitates higher heterologous expression. Nucleosome Density additionally trended lower in the high-expression group (p = 5.40×10⁻², Δ = –0.35). Collectively, the strongest signals arise from distinct feature categories, like chromatin structure, neighborhood activity, and nucleosome occupancy, indicating that position effects result from integrated contributions across multiple regulatory layers.

Next, we assessed the individual predictive contribution and potential collinearity among engineered features. Single-feature linear regressions revealed that while “Boundary Distance” and “Genome Compact Score” exhibited the highest individual predictive explanatory power, yet all features yielded low R² values, indicating that no single parameter can account for the expression variance (Supplementary Fig. S2). Pairwise correlation analysis further revealed minimal multicollinearity across features, except for the expected correlation between H3K4me1 and H3K4me2 enrichment (Supplementary Fig. S3). These results collectively support that position effects are inherently multi-factorial, which can only be accurately modelled by integrating multiple genomic layers.

### Model performance and inference

Given the limited size of our dataset (N=98), we adopted classical gradient-boosting methods, including XGBoost, CatBoost, LightGBM, and hyper-parameter search via the AutoGluon package (Fig. 2A). Due to the small size and heavy-tailed distribution of fluorescence intensity, we employed SPCC as the primary evaluation metric, with PCC, R^2^, and MSE being reported as supportive calibration metrics. Across model configurations, the SPCC values consistently converged between 0.4 and 0.6, with a peak density between 0.45∼0.55 on the held-out test set (Fig. 2B). The six best configurations show consistent generally monotonic relationships on the test set (Supplementary Fig. S4), supporting model performance is reproducible across different learners and settings. Notably, out-of-fold errors were lower than those observed on held-out test (Supplementary Fig. S5), indicating mild overfitting, which is a common occurrence given the small tabular dataset. To confirm that the model’s predictive capability stems from genuine biological signals rather than statistical noises, we performed a target permutation test with label-scrambling. While the model trained on the whole dataset (N=98) demonstrated strong predictive power (Overall SPCC = 0.693), the performance collapsed to near-random guessing (SPCC = 0.102) when expression labels were randomly shuffled (Supplementary Fig. S6).

**Figure 2.**
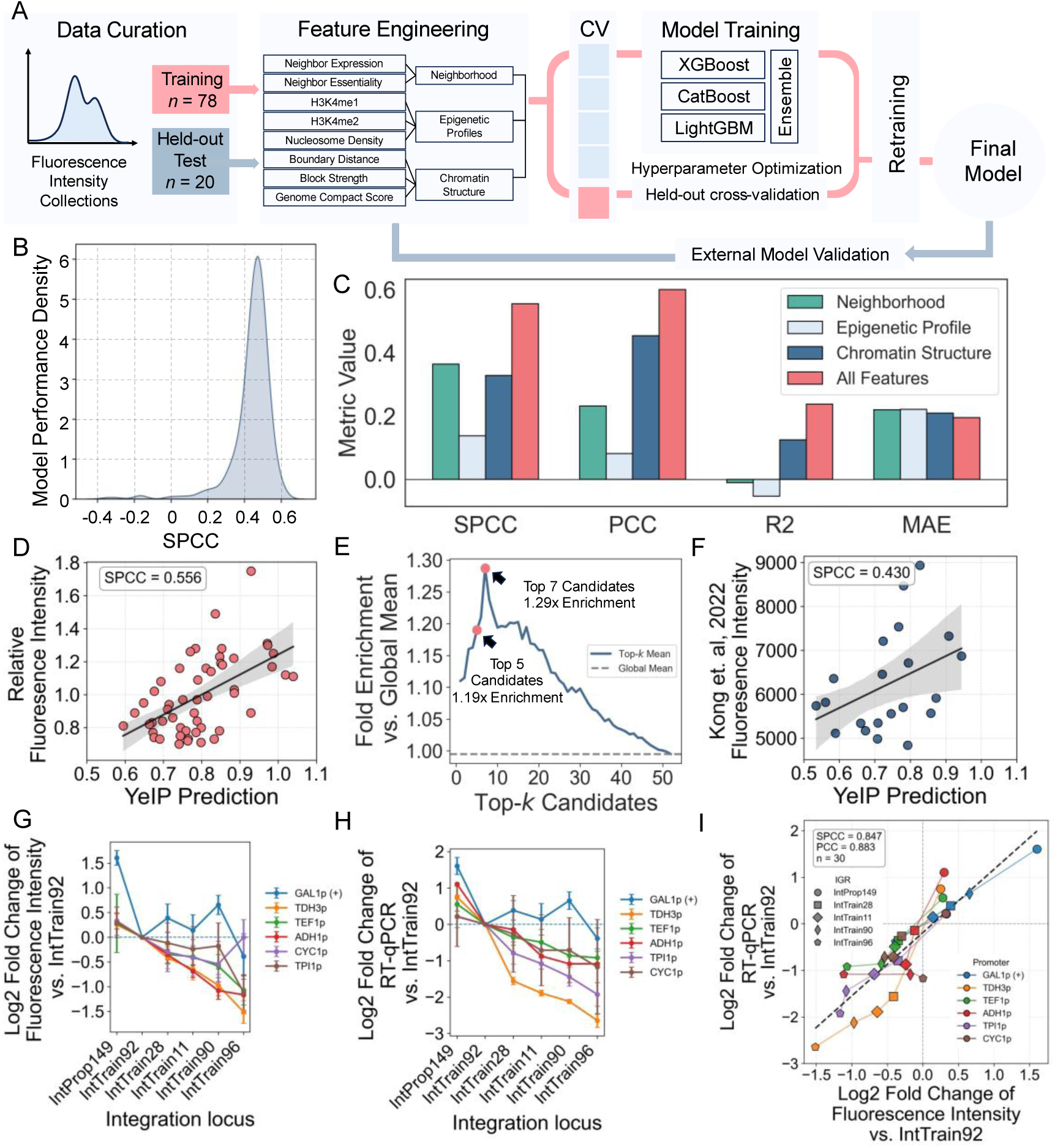
YeIP training, validation, and interpretation. (A) Schematic overview of YeIP machine learning training pipeline. (B) Performance distribution of models generated during the training process. The density plot displays the Spearman correlation coefficients (SPCC) of trained models on the test set. (C) Feature ablation analysis evaluating the contribution of distinct genomic feature groups. Performance metrics (SPCC, PCC, R2, MAE) are compared for models trained using only Transcriptional Neighborhood (teal), Epigenetic Profile (light blue), Chromatin Structure (dark blue), or All Features combined (red). (D) Experimental validation of YeIP on 52 unseen integration sites (Supplementary Table S2). (E) Top-*k* enrichment analysis validating the model’s ranking capability, where curve displays the fold enrichment of the mean fluorescence intensity for the top candidates relative to the global mean of the 52 unseen integration sites. (F) External validation comparing YeIP predictions with experimentally measured fluorescence intensities from an independent dataset (Kong et al., 2022). (G) Normalized fluorescence intensity relative to reference site IntTrain92 of 6 different promoters integrated into 6 representative IGRs. (H) Relative mRNA abundance relative to IntTrain92 of promoter-IGR combinations quantified by RT-qPCR. (I) Scatter plot showing the correlation between RT-qPCR and fluorescence intensity across the tested promoter-IGR combinations.

We evaluated the contribution of each feature class using ablation analysis (Fig. 2C), where the independent predictive power of each biological layer was evaluated by training stand-alone models on specific feature subsets. We grouped features as “Neighborhood”, “Epigenetic profile”, “Chromatin profile”. Group settings followed the feature definition mentioned in the former section. The full ten-feature model achieved the highest performance (SPCC=0.556), demonstrating complementarity contributions across feature categories (Fig. 2D). Transcriptional Neighborhood alone exhibited the strongest single-group predictive power (SPCC=0.372). Chromatin structural features contributed moderately (SPCC=0.330). Epigenetic marks alone showed limited predictive value (SPCC=0.144). These results reinforce that positional effects emerge from distributed multi-layer genomic determinants, and that multi-feature integration is necessary to preserve rank fidelity (Supplementary Fig. S7).

We selected the model with the highest SPCC for YeIP and validated it using an independent experimental batch of 52 new integration sites (hereafter referred to as “IntProp” loci; Supplementary Table S2) that were excluded from the training phase. YeIP accurately enriched for high-performing loci (SPCC=0.556, Fig. 2C), demonstrating strong rank fidelity of YeIP. To quantify the practical utility of YeIP for prioritizing high-performing sites, we performed a Top-*k* enrichment analysis (Fig. 2E). This metric evaluates the fold-increase in mean fluorescence intensity of the top k candidates predicted by the model relative to the global average of the 52 unseen integration sites. The analysis revealed a robust enrichment signal, peaking at the top 7 candidates with a 1.29-fold increase over the global mean. Note that IntTrain92 is a relatively high expression reference IGR, the curve still remains consistently above the baseline, indicating that the model effectively concentrates high-expression IGRs at the top of its ranking list. SHapley Additive exPlanations (SHAP) analysis further confirmed that the model predictions are driven by biologically coherent features consistent with established determinants of transcriptional output, including chromatin accessibility and neighborhood transcriptional activity (Supplementary Fig. S8). We then tested YeIP against an external, independent published dataset (19) and observed a positive rank correlation (SPCC=0.430, Fig. 2F). Although this correlation was more modest than that observed in our independent experimental validation set, such difference may result from the variations in reporter design, strain background, and measurement protocol between datasets. This independent validation supports that YeIP captures generalizable rules underlying position effects, instead of overfitting to the specific training data of this study.

To assess the universality of these position effects, we extended our testing to 6 distinct promoters with varying strength, including the inducible *GAL1p*, across 6 representative IGRs. The relative expression ranking of the IGRs remained highly consistent across *TDH3p, TEF1p, ADH1p, CYC1p,* and *TPI1p* contexts (Fig. 2G). Notably, *GAL1p* showed a modest deviation from the trends observed for the constitutive promoters, while IntProp149 remained among the top-performing loci under *GAL1p* and exceeded the reference integration site in fluorescence intensity. We observed a small deviation for the *CYC1p*– IntTrain96 combination, possibly because the relatively low expression level of *CYC1p* makes the measurement more susceptible to variation under low-signal conditions, while also suggesting that post-transcriptional effects may contribute to the difference between mRNA and fluorescence readouts. Detailed characterization of the promoter and IGR combination matrix (Supplementary Fig. S9) further clarified that while promoter identity dominated the absolute magnitude of expression, IGR exerted a reproducible, scalar modulation on top of this baseline. This orthogonality was consistent across both phenotypic and transcriptional readouts. RT-qPCR analysis of *TDH3p, TEF1p, ADH1p, CYC1p, TPI1p,* and *GAL1p* combinations confirmed a strong monotonic association (SPCC = 0.847, N=30) between mRNA abundance and fluorescence intensity (Fig. 2H, 2I, and Supplementary Fig. S10). Notably, although the relative locus ranking was generally preserved after normalization to IntTrain92 within each promoter context, the magnitude of locus-dependent modulation differed among promoters. Stronger promoters such as *TDH3p* showed a broader dynamic range across loci, whereas weaker promoters such as *CYC1p* exhibited a more compressed response. To assess environmental and genetic robustness, we evaluated IGR expression across different carbon sources. The relative locus ranking remained highly conserved across fermentable sugars (dextrose, galactose, and raffinose), whereas non-fermentable glycerol modulated this ordering (Supplementary Fig. S11). Overall, these results confirmed that IGR-dependent regulation is a persistent genomic feature across diverse carbon sources.

Together, these data suggest that, within the tested construct panel and growth conditions, YeIP captures a reproducible locus-dependent ranking that is primarily reflected at the mRNA level.

### Genome-wide inference of expression hot spots and cold spots

To further establish a genome-scale resource for rational design of yeast expression systems, we then employed YeIP to predict expression potential for 589 IGRs (Supplementary Table S5). This final list of 589 IGRs was refined from an initial pool of a genome-wide 6292 IGRs by applying stricter exclusion criteria for centromeric (<0.5 kb) and telomeric (<5 kb) regions to ensure genetic stability, alongside the >800 bp length filter (Supplementary Fig. S1). It should be noted that YeIP reports a dimensionless score calibrated to the training distribution that is intended for ranking and top-K enrichment (Fig. 2E), not for absolute expression estimates. Genome-wide summaries therefore show predicted scores rather than fold-changes. To investigate whether expression potential is dominated by proximity to large chromosomal landmarks, we mapped both the experimental training data and the genome-wide YeIP predictions against their distances to the nearest centromeres and telomeres (Fig. 3A and 3B). The lack of a simple linear gradient suggests that expression potential is not solely determined by the physical distance to these landmarks, but rather by the complex local chromatin features captured by YeIP. This confirms that the chromosomal arms provide a broad and tunable landscape, populated with numerous candidate sites spanning a wide dynamic range.

**Figure 3.**
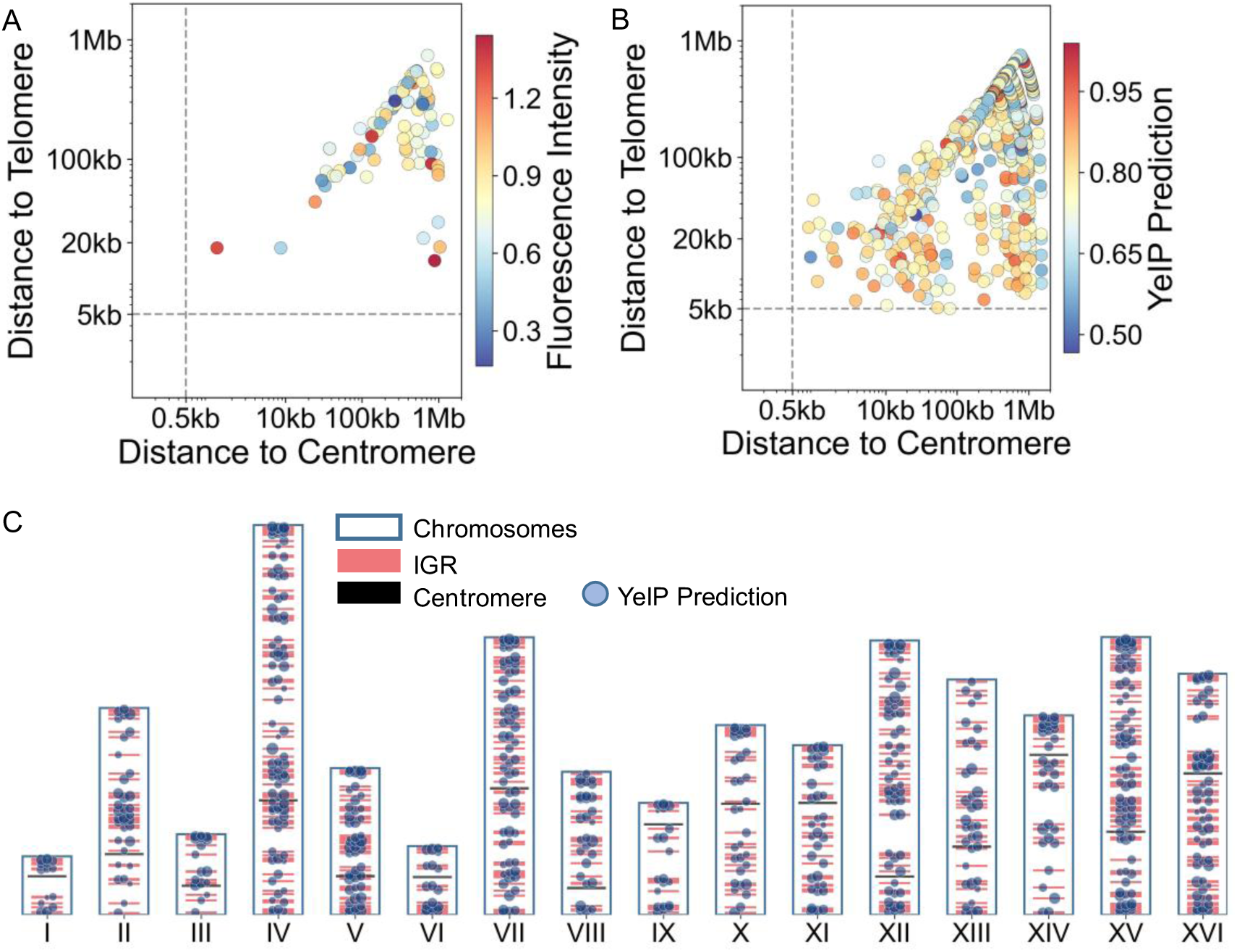
Genome-wide atlas and spatial distribution of expression potential for IGRs in *S. cerevisiae*. (A, B) Spatial analysis of expression potential relative to chromosomal landmarks. Scatter plots display the relationship between the distance to the nearest centromere and telomere for (A) the experimental training dataset and (B) the genome-wide YeIP predictions. Points are colored by (A) measured fluorescence intensity or (B) predicted YeIP score. (C) Schematic overview of predicted expression potential across all 16 chromosomes. Each blue circle represents an engineering qualified IGR, with the circle’s diameter proportional to its YeIP prediction score. Red marks indicate the relative location of the IGRs, and black blocks denote the centromeres (Supplementary Table S5).

The genome-wide prediction map is visualized in Fig. 3C, where the size of each circle is proportional to the predicted expression strength of its corresponding IGR. The genome-wide predictions followed an approximately normal distribution (mean ≈ 0.73), closely matching that of the experimental dataset and indicating the model is well-calibrated (Supplementary Fig. S12). Such alignment suggests that YeIP accurately extrapolates beyond the training set, capturing the natural variability of chromosomal regulatory environments. To explore the spatial organization of position effects, we computationally aggregated predictions into 50-kb sliding windows across the genome (Supplementary Fig. S13), revealed that contiguous chromosomal domains exhibit coordinated high or low predicted expression levels, reflecting the underlying intermediate-scale structure of yeast chromatin. We then compiled a ranked catalog of the top ten high-activity “hot regions” and low-activity “cold regions” genomic domains defined at this 50-kb window scale (Supplementary Table S6), typically encompassing 3∼7 IGRs per regions.

To validate this lower boundary, we experimentally tested 16 additionally predicted cold spots, all of which exhibited uniformly strong repression (Supplementary Fig. S14). Importantly, while regional averaging smooths predicted scores, individual IGRs exhibit a much broader ∼10-fold dynamic range (Supplementary Fig. S15). Together, these results confirm that chromosomal position effects provide a robust, predictable regulatory window for precise gene expression fine-tuning.

### Tuning the lycopene biosynthesis pathway via strategic site selection

To demonstrate the practical utility of the genome-wide expression atlas, we further applied it to the optimization of a heterologous pathway. As a model demonstration, we chose the three-enzyme lycopene biosynthesis pathway, consisting of CrtE (geranylgeranyl pyrophosphate synthase), CrtB (phytoene synthase), and CrtI (phytoene desaturase) (Fig. 4A). Lycopene is a vibrant red pigment, allowing for a direct visual assessment of pathway productivity via colony color. We then constructed a combinatorial library in which each gene was integrated into distinct IGRs with relatively high, medium, and low reporter fluorescence levels characterized by fluorescence intensity measurements, thereby generating strains with diverse tri-gene expression configurations.

**Figure 4.**
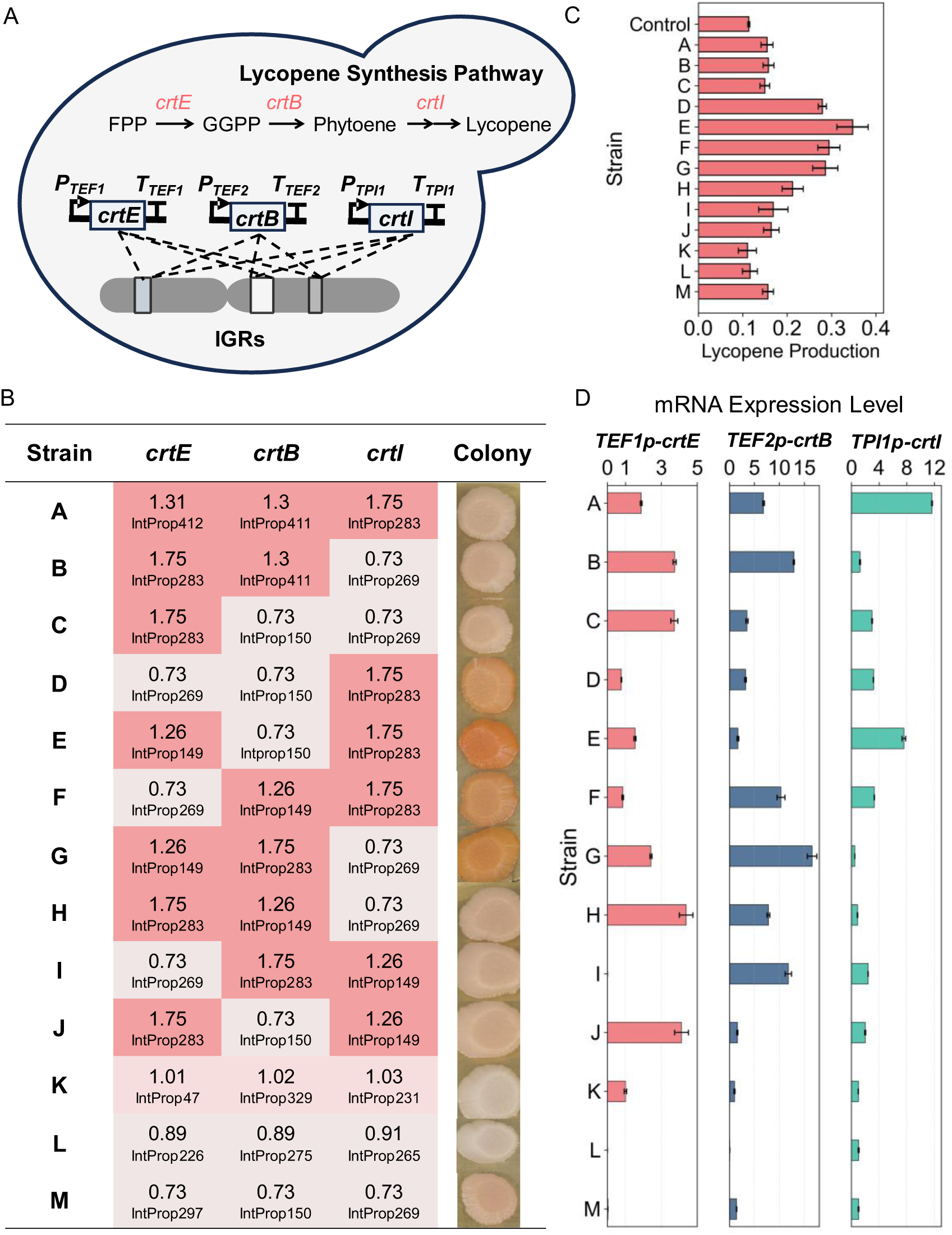
Combinatorial tuning of the lycopene biosynthesis pathway with IGRs. (A) Schematic of the experimental design for pathway optimization. (B) Library design showing the combinatorial assignment of the three pathway modules to genomic loci. Values indicate the experimentally measured relative fluorescence intensity of each locus (Supplementary Table S2). Representative colony images illustrate the resulting pigmentation spectrum. (C) Quantification of lycopene production for the engineered strains by OD_470_ nm/OD_600_ nm absorbance with a strain lacking the lycopene biosynthesis pathway as control. (D) RT–qPCR measurements of *crtE, crtB*, and *crtI* across the same strain panel. Values represent fold changes (2^−ΔΔCt^) relative to the calibrator strain (Strain K) for each target gene after normalization to *ALG9*; values close to zero indicate strong downregulation relative to Strain K rather than absence of expression.

The library displayed a broad spectrum of pigmentation phenotypes (Fig. 4B). Strains with very weak or cream-colored colonies were associated with either globally low expression or strongly imbalanced expression patterns, indicating that certain IGR combinations restrict flux at early or late steps. Strains with pale pink or light orange coloration reflected partially matched expression levels. A subset of combinations produced deep orange-red colonies, corresponded to high lycopene accumulation. Consistently, lycopene absorbance measurements of cell extracts (Fig. 4C) confirmed that strains with deeper pigmentation exhibited higher lycopene content, supporting the visual phenotype classification.

These high-performing strains represent configurations in which *crtE, crtB,* and *crtI* transcription levels are proportionally balanced to support efficient conversion of the native farnesyl pyrophosphate (FPP) pool. The observed phenotypes are congruent with known bottlenecks in carotenoid pathways (21,36,37). For instance, the non-pigmented phenotype of strain C, integrated the pathway genes at IGRs with previously measured relative fluorescence values of 1.75 for *crtE*, 0.73 for both *crtB* and *crtI*, supports findings that excessive upstream CrtE activity may create a metabolic burden by draining the FPP precursor pool. Similarly, strain B, integrated the pathway genes at IGRs with measured relative fluorescence values of 1.75 for *crtE*, 1.3 for *crtB*, and 0.73 for *crtI*, suggests that the desaturation step catalyzed by CrtI is rate-limiting when substrate is plentiful, leading to the accumulation of a colorless intermediate (38,39). Notably, we failed to achieve maximal lycopene production on strain A by uniformly increasing the expression of all three enzymes simultaneously (i.e., by integrating all genes into high-reporter fluorescence levels IGRs). Instead, the most productive variant, strain E, emerged from a specific, rationally balanced expression profile, where the pathway genes are integrated at measured relative fluorescence values of 1.26 for *crtE* expression, 0.73 for *crtB* expression, and 1.75 for *crtI* expression. This surprising configuration indicates that while alleviating pathway bottleneck (CrtI) is critical, excessive levels of the intermediate enzyme (CrtB) are unnecessary and potentially detrimental to overall metabolic flux.

Importantly, RT-qPCR measurements of the pathway genes (*crtE, crtB,* and *crtI*) across the engineered strains confirmed that these phenotypic differences were driven by precise, site-specific transcriptional modulation. Transcript abundance strongly correlated with the measured relative fluorescence values of the integration loci (Fig. 4D and Supplementary Fig. S16), providing direct evidence that IGR tuning achieves rational stoichiometric balancing at the mRNA level.

These results demonstrate that the chromosomal integration sites can function as effective regulatory elements to titrate multi-gene expression and achieve stoichiometric balance. The characterized IGRs portfolio with varied expression levels function as a powerful genomic rheostat, enabling the practical discovery of optimal expression ratios and shifting pathway optimization from empirical screening toward rational, model-informed design.

## Discussion

This study establishes a data-driven framework for predictable genome engineering in *S. cerevisiae*. We developed the YeIP model to predict gene expression level directly from genomic context and constructed a genome-wide atlas identifying expression hotspots and coldspots. By integrating this atlas into metabolic pathway design, we demonstrated that chromosomal integration site can act as an effective regulatory layer for tuning pathway transcriptional level and potentially balancing the metabolic flux. These findings reposition chromosomal context from a passive background variable into a programmable element of synthetic biology, enabling rational, location-informed control over heterologous expression.

Given the limited size of our experimental dataset, we trained YeIP with an ensemble of gradient boosting methods, which is optimal for capturing non-linear biological interactions in small tabular data where deep learning is prone to overfitting. While YeIP provides a robust framework for positional tuning, fully harnessing its potential requires further defining its biological boundary conditions. Currently, we train and validate the model primarily using transcriptomic and chromatin features derived from haploid cells under standard YPD growth conditions (14,23–27). Encouragingly, our targeted evaluations reveal that relative locus rankings are highly conserved across alternative fermentable carbon sources (e.g., galactose and raffinose). However, strong physiological shifts, such as growth in non-fermentable glycerol, can modulate this ordering (Supplementary Fig. S11). An additional frontier is the potential interplay between specific IGRs and promoter elements. Although promoter generally dominate absolute expression (40), local crosstalk between integrated sequences and neighboring chromatin environments cannot be excluded. Systematically profiling large-scale promoter-IGR matrices will clarify whether such interactions introduce predictable biases or stochastic deviations, for further refining the precision of locus-based expression tuning.

Crucially, YeIP-guided integration offers a complementary strategy to traditional promoter engineering. While promoters provide “coarse” control with log-scale changes, IGR tuning offers construct-preserving “fine” control with linear-scale, which is essential for alleviating metabolic burden and achieving stoichiometric balance. Beyond metabolic engineering, this framework has broad implications. For genetic circuits, it allows spatial tuning of repressor/activator ratios without altering promoter logic, ultimately advancing the standardization of yeast cell factories. By transforming undefined intergenic regions into characterized synthetic biological parts, YeIP is expected to facilitate the automated assembly of multi-gene systems in biofoundries, turning the genome into a standardized chassis.

In conclusion, the YeIP model and its accompanying genome-wide atlas provide a robust, data-driven foundation for rational metabolic pathway engineering in yeast. By transforming position effects into a quantifiable and designable parameter, our work moves the field beyond empirical tuning toward predictive genome-scale design, offering a new dimension of control for synthetic biology and industrial biotechnology.

## Data Availability

The gRNA sequences for experimentally characterized IGRs, the genome-wide inference results, and the complete genome-scale feature matrix are available in the supplementary data. To facilitate reproducibility, the source code for feature extraction, model training, and inference is publicly available on GitHub at https://github.com/daftpunksss/YeIP.

## Acknowledgments

We thank Professor Shuobo Shi from Beijing University of Chemical Technology for sharing yeast strains with *mCherry* expression cassette integrated at different loci. We also would like to thank iBioFoundry and Core Facility at ZJU-Hangzhou Global Scientific and Technological Innovation Center for analytical support.

## Author contributions

H.H. developed YeIP and drafted the manuscript. Y.C. and J.W. performed all the experiments. H.H. and Y.C. analyzed the data. C.D., Q.Z., and J.L. conceived the study and revised the manuscript. All authors approved the manuscript.

## Supplementary Materials

Supplementary Data are available.

