## Supplementary material for "Exploring Chromosomal Position Effects for Predictable Tuning of Metabolic Pathways in Yeast": SI

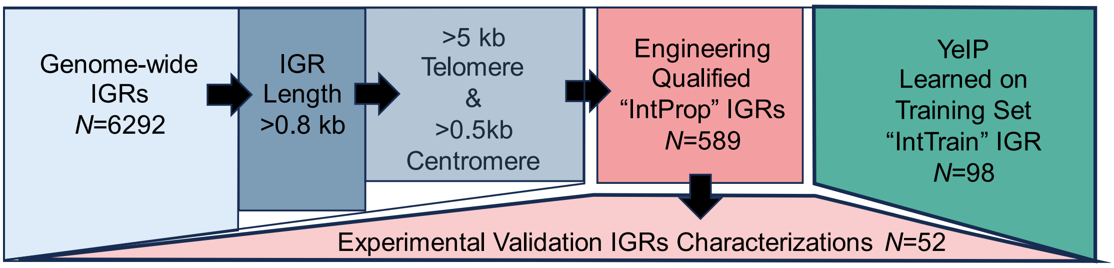


**Figure S1. Data curation and workflow for genome-wide IGR selection.** Flowchart illustrating the stepwise filtering process used to generate the datasets. The study started with all 6292 annotated IGRs. The Training Set (N=98) was selected based on high experimental integration efficiency (>75%) and included a range of lengths to train the YeIP model. For the Genome-wide Prediction, IGRs were filtered to exclude unstable centromeric/telomeric regions and short sequences (<800 bp), resulting in 589 engineering qualified IGRs recommended for synthetic biology applications. The Validation Set (N=52) consists of independent IGRs used to test model performance on unseen data.

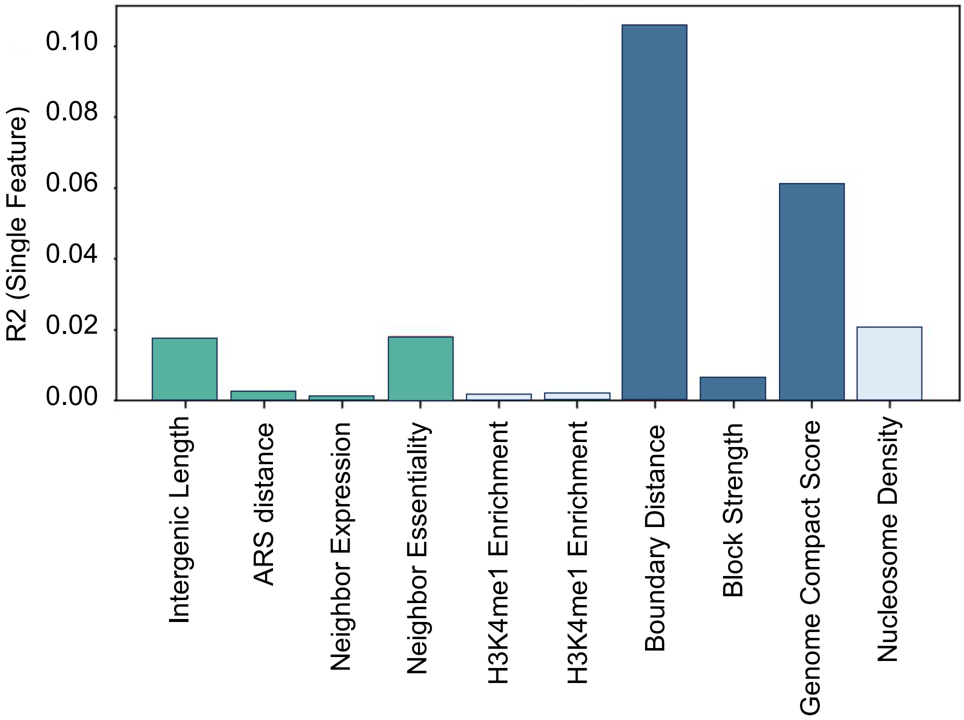


**Figure S2. Explained variance by individual genomic features.** To evaluate the individual predictive power of each engineered feature, a series of single-feature Ordinary Least Squares (OLS) linear regression models was trained. Each model used only one feature (x-axis) to predict the measured fluorescence intensity. The y-axis shows the coefficient of determination ($R^2$) for each model.

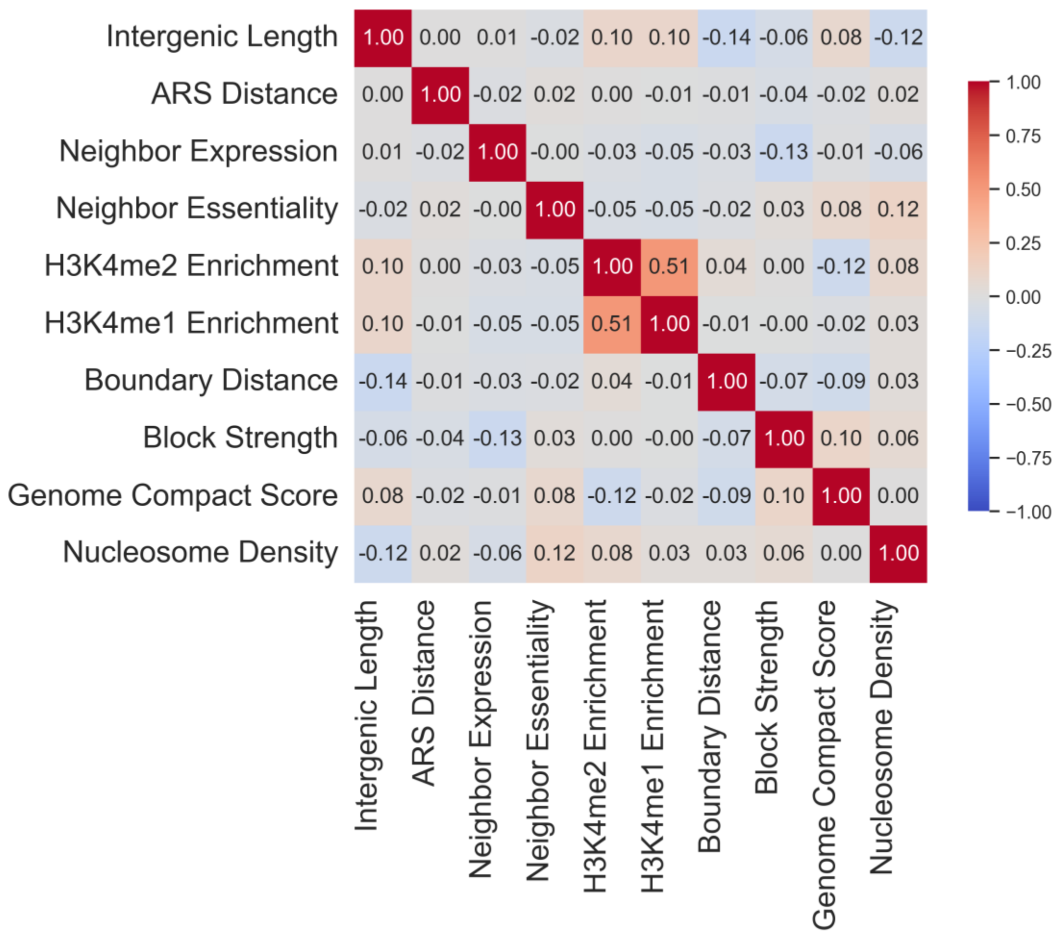


**Figure S3. Correlation matrix of engineered features.** Heatmap showing the Pearson correlation coefficients between all pairs of engineered features used in YeIP. The color scale indicates the strength of the correlation, from strongly positive (red, +1.00) to strongly negative (blue, -1.00).

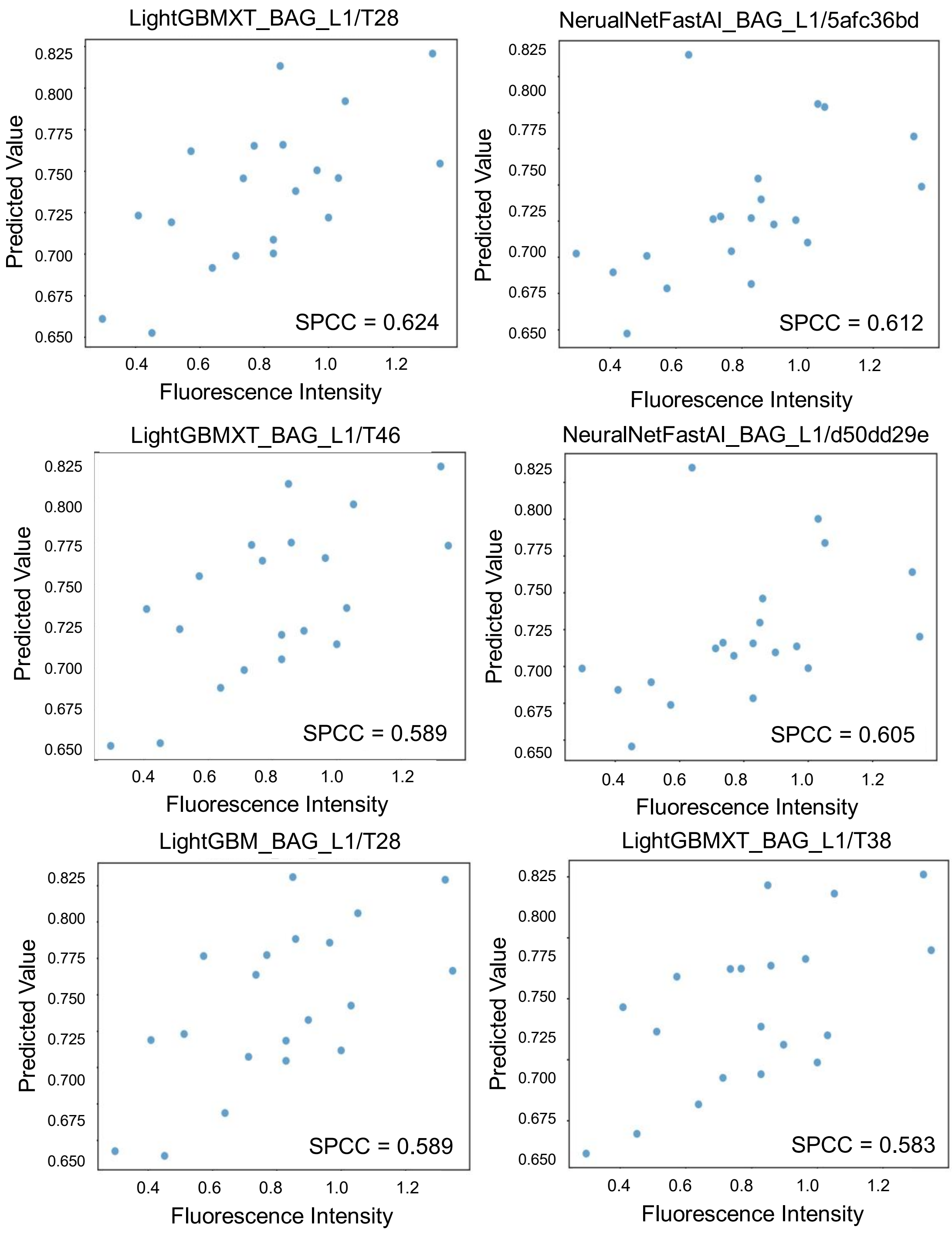


**Figure S4. Performance of the six best-performing individual models on the held-out test set.** Each scatter plot compares the predicted expression value (y-axis) against the experimentally measured fluorescence intensity (x-axis) for one of the top-ranked models generated during the AutoGluon training process. The title of each plot indicates the specific model architecture and training run. Predicted Value represents the specific model output.

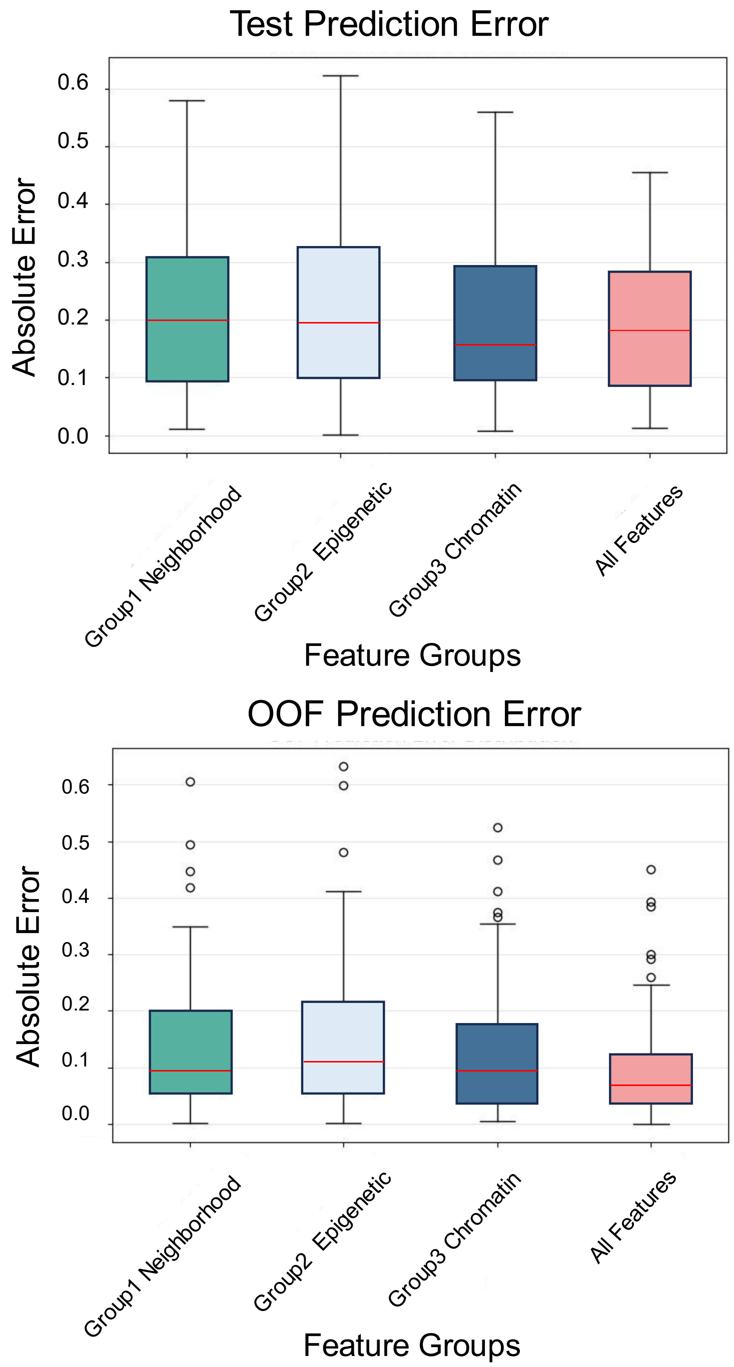


**Figure S5. Distribution of absolute prediction errors for ablation study models.** Box plots comparing the absolute prediction errors from models trained on distinct feature subsets as part of the ablation study. The feature groups are: Neighborhood features (Group1), epigenetic features (Group2), chromatin group (Group3), and All Features. (Top) Distribution of absolute errors for out-of-fold (OOF) predictions across the entire training dataset. (Bottom) Distribution of absolute errors for the same models on the held-out test set.

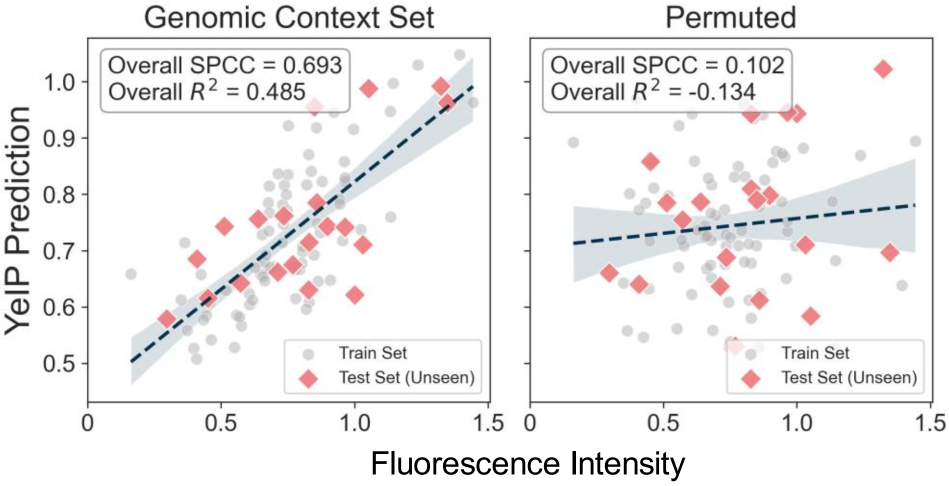


**Figure S6. Permutation control for genomic-context features.** YeIP predictions versus measured expression for models trained with the full genomic-context feature set (left) or with the same features randomly permuted across loci (right). Grey circles indicate training samples and red diamonds indicate held-out test samples. Dashed lines show linear fits with 95% confidence bands. Overall Spearman correlation (SPCC) and $R^2$ are reported in each panel.

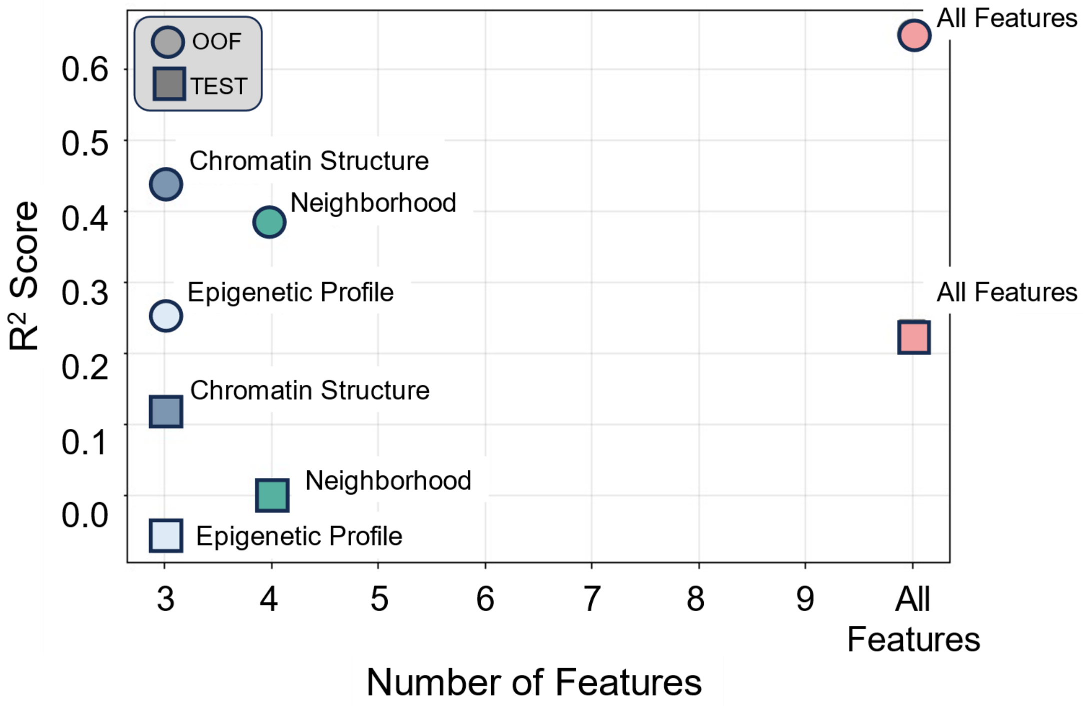


**Figure S7. Model performance as a function of feature count.** Scatter plot summarizing the results of the feature ablation study. The x-axis represents the number of features used to train each model, and the y-axis represents the resulting $R^2$ score. Performance is shown for both out-of-fold (OOF) predictions on the training data (circles) and predictions on the held-out test set (squares).

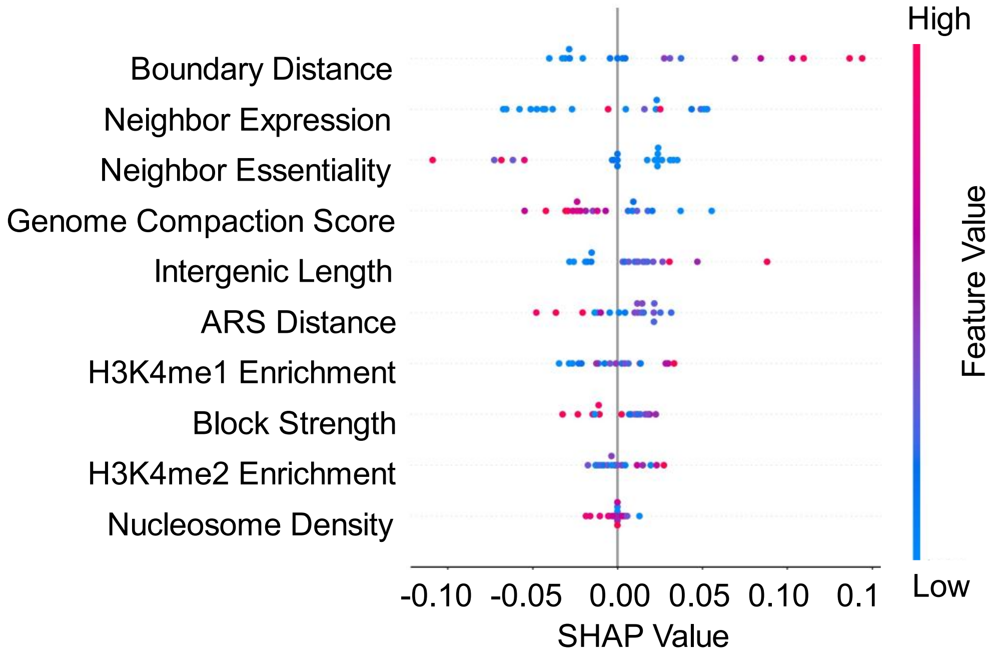


**Figure S8. SHAP summary plot for the final YeIP.** The plot provides a detailed interpretation of the model's predictions by showing the impact of each feature. Features (y-axis) are ranked by their global importance, calculated as the mean absolute SHAP value. Each point on a feature's row represents an individual IGR from the dataset. The position of the point on the x-axis indicates the SHAP value, which represents the impact of that feature on the model's output for that specific IGR; positive values push the prediction higher, while negative values push it lower. The color of each point corresponds to the feature's value for that IGR, ranging from low (blue) to high (red).

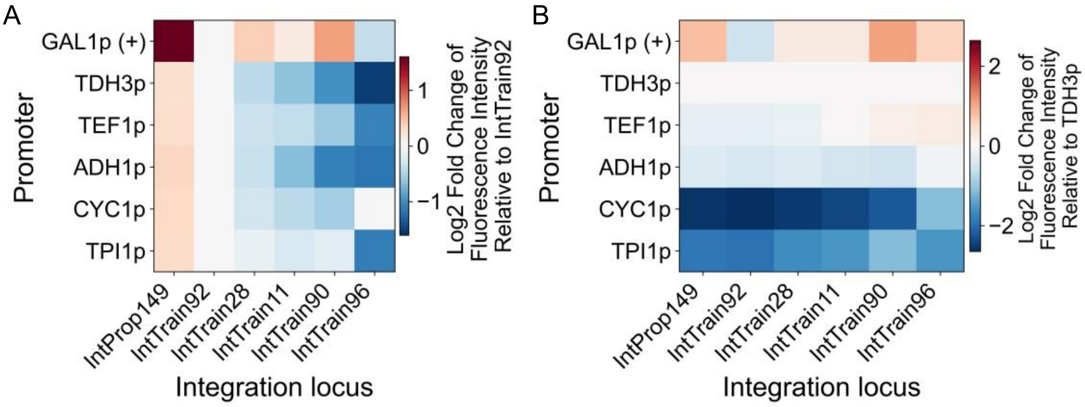


**Figure S9. Promoter–locus fluorescence intensity in the promoter × integration-site panel.** (A) Fluorescence measurements shown as log2 fold-change relative to IntTrain92. (B) Fluorescence measurements shown as log2 fold-change relative to *TDH3p* within each locus. Promoters and integration loci are indicated on the axes; color scales are shown beside each heatmap.

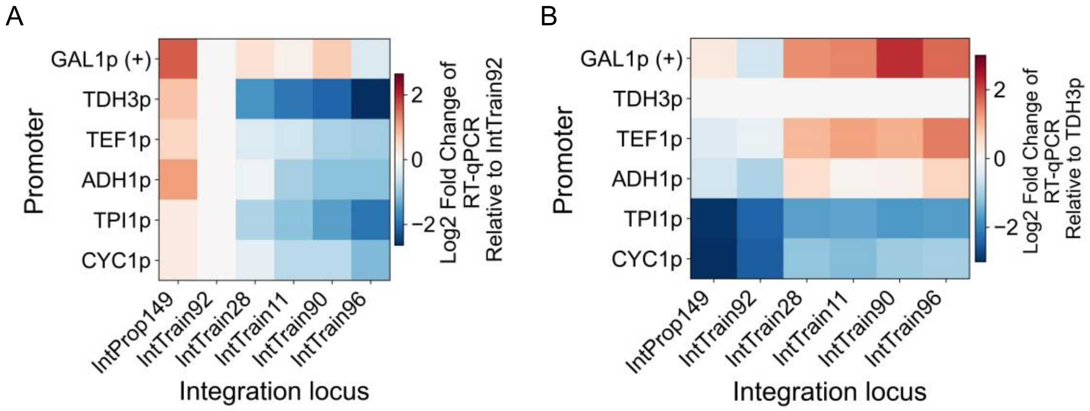


**Figure S10. Promoter–locus RT-qPCR in the promoter × integration-site panel.** (A) RT–qPCR results shown as log2 fold-change relative to IntTrain92. (B) RT–qPCR results shown as log2 fold-change relative to *TDH3p* within each locus. (C) Fluorescence measurements shown as log2 fold-change relative to IntTrain92. (D) Fluorescence measurements shown as log2 fold-change relative to *TDH3p* within each locus. Promoters and integration loci are indicated on the axes; color scales are shown beside each heatmap.

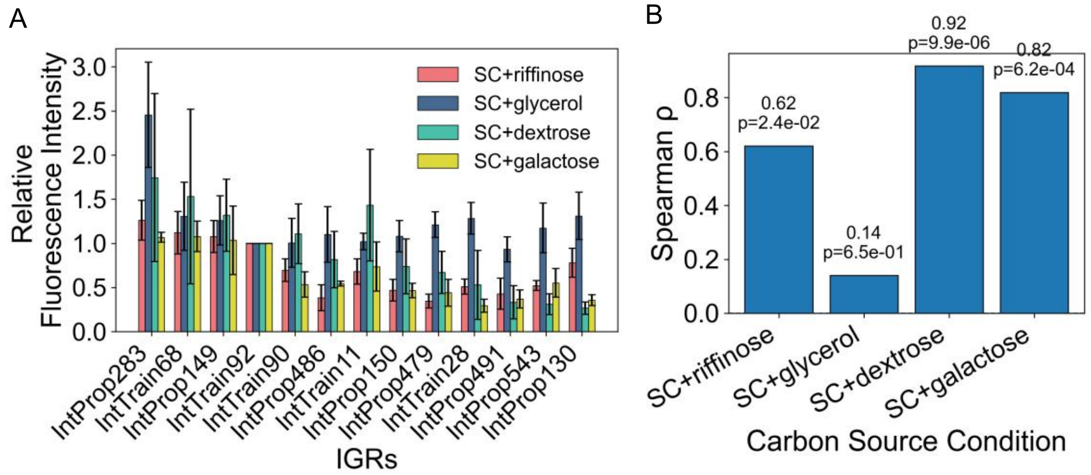


**Figure S11. Position-effect measurements across carbon source conditions.** (A) Relative fluorescence intensity measured for the indicated integration loci under four SC media with different carbon sources (raffinose, glycerol, dextrose, and galactose). Loci are ordered by their expression level under the standard SCD condition. Fluorescence intensity is normalized with IntTrain92 (*TDH3p*, SCD medium). (B) Spearman rank correlation coefficient (ρ) between the locus ranking observed in each carbon source condition and that measured under SCD.

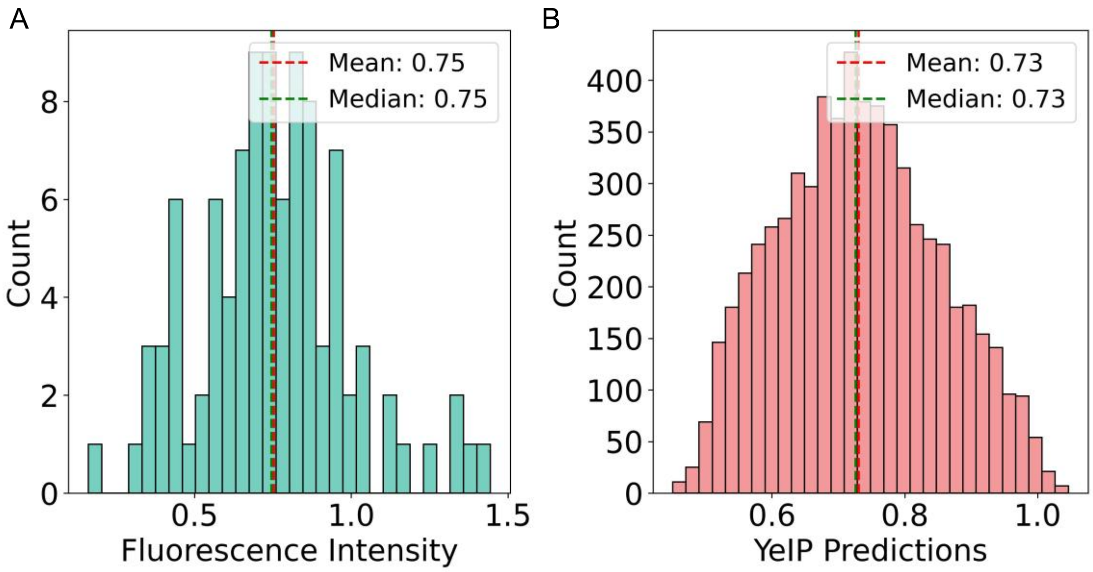


**Figure S12. Comparison of experimental and predicted expression distributions.** (A) Histogram of the experimentally measured fluorescence intensity for the 98 IGRs that comprise the training dataset. (B) Histogram of the YeIP's predicted expression values for all 589 long IGRs across the genome.

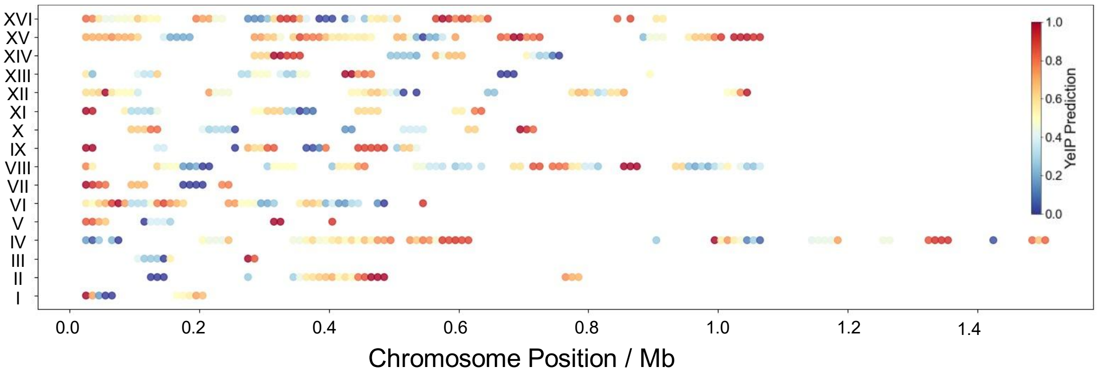


**Figure S13. Linear map of YeIP predictions across all 16 *Saccharomyces cerevisiae* chromosomes.** Each point represents a 50-kb scanning window (IGR), positioned by its chromosomal coordinate (Mb) on the x-axis. The y-axis denotes the 16 yeast chromosomes (Roman numerals I to XVI). The predicted expression potential for each IGR is indicated by the color scale, ranging from low (blue, ~0.0) to high (red, 1.0).

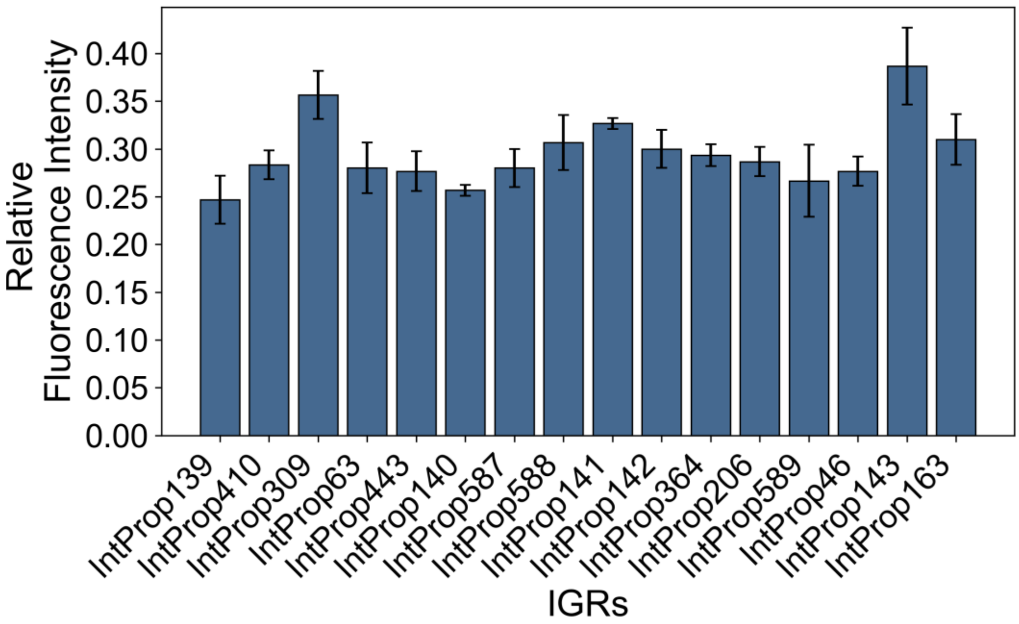


**Figure S14. Measurements of additional low-activity IGR.** Relative fluorescence intensity measured for the indicated set of integration loci selected from the low predicted-expression range. Fluorescence intensity is normalized with IntTrain92 (*TDH3p*, SCD medium).

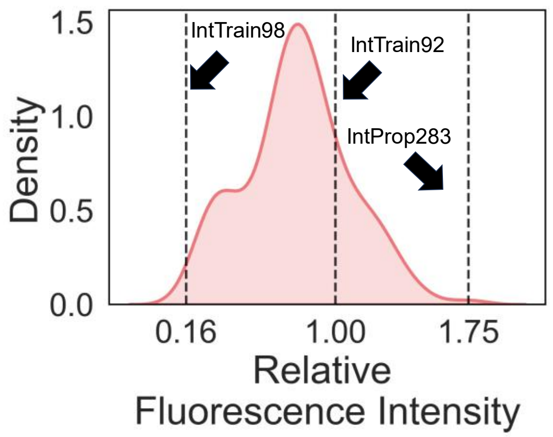


**Figure S15. Distribution of measured fluorescence intensities normalized to IntTrain92 (*TDH3p*, SCD medium) across all experimentally characterized IGRs in this article.** The three vertical dashed lines denote the weakest expression site (IntTrain98, ~0.16), the reference site (IntTrain92, 1.00), and the strongest expression site (IntProp283, ~1.75).

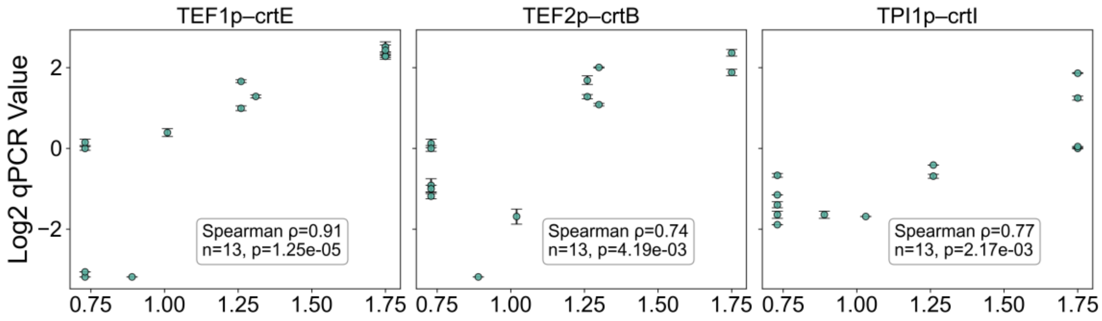


**Figure S16. Rank Correlations between IGR fluorescence intensity and module-level RT–qPCR in the lycopene strain panel.** Scatter plots show, for each strain (n = 13), the relationship between IGR fluorescence intensity and the corresponding log2 fold change RT–qPCR values normalized with *ALG9* and using Strain D as the calibrator for the three pathway modules: *TEF1p–crtE*, *TEF2p–crtB*, and *TPI1p–crtI*. Spearman rank correlation coefficient (ρ) and p-values are indicated in each panel.
